# Retinal degeneration: Multilevel protection of photoreceptor and ganglion cell viability and function with the novel PKG inhibitor CN238

**DOI:** 10.1101/2021.08.05.455191

**Authors:** Arianna Tolone, Wadood Haq, Alexandra Fachinger, Andreas Rentsch, Friedrich W. Herberg, Frank Schwede, François Paquet-Durand

## Abstract

Hereditary retinal degeneration (RD) is often associated with excessive cGMP-signaling in photoreceptors. Previous research has shown that inhibition of cGMP-dependent protein kinase G (PKG) can slow down the loss of photoreceptors in different RD animal models. In this study, we identified a novel PKG inhibitor, the cGMP analogue CN238, with strong protective effects on photoreceptors in retinal degeneration *rd1* and *rd10* mutant mice. In long-term organotypic retinal explants, CN238 preserved *rd1* and *rd10* photoreceptor viability and function. Surprisingly, in explanted retinae CN238 also protected retinal ganglion cells from axotomy induced retrograde degeneration and preserved their functionality. Together, these results confirm the strong neuroprotective capacity of PKG inhibitors for both photoreceptors and retinal ganglion cells, thereby significantly broadening their potential applications for the treatment of retinal diseases and possibly neurodegenerative diseases in general.

## Introduction

Photoreceptor degeneration is a hallmark of retinal degenerative diseases (RD), a group of retinal dystrophies characterized by primary dysfunction and degeneration of photoreceptor cells, leading to visual loss and eventually to blindness (Berger, Kloeckener-Gruissem, and Neidhardt (2010). Because of the genetic heterogeneity of RD-type diseases, no effective therapies are currently available (Sahel, Marazova, and Audo 2014). While the mechanisms causing photoreceptor degeneration are far from being fully understood, high levels of cyclic guanosine monophosphate (cGMP) are known to trigger non-apoptotic photoreceptor cell death in many RD disease models (Arango-Gonzalez et al. 2014; Power et al. 2020). Two commonly used models for studying RD are *rd1* and *rd10* mice, which are characterized by a nonsense (*rd1*) and a missense (*rd10*) mutation in the gene encoding for the β-subunit of rod phosphodiesterase (PDE) 6 (Han et al. 2013). Lack of PDE6 activity leads to an accumulation of cGMP in photoreceptors (Farber and Lolley 1974; Paquet-Durand et al. 2009) and emerging RD neuroprotection strategies include targeting pathways downstream of cGMP (Arango-Gonzalez et al. 2014; Power et al. 2020).

The prototypic cellular target of cGMP is protein kinase G (PKG), a serine/threonine kinase that exists as a homodimer of two subunits, each consisting of an N-terminal dimerization domain, an auto-inhibitory sequence, two cGMP binding sites, and a C-terminal kinase domain. Three isoforms of PKG have been identified in mammals: PKG1α, PKG1β and PKG2. When cGMP binds to PKG, it induces the release of the C-terminal catalytic domain from the auto-inhibitory sequence, activating PKG (Kim et al. 2016). cGMP-signaling activates PKG, which when overactivated is likely to play a key role in triggering cell death (Browning 2008; Power et al. 2020; Canals et al. 2003; Canzoniero et al. 2006; Fallahian et al. 2011; Leung et al. 2010). In the retina, exceedingly high cGMP levels, as well as strong PKG activation were causally linked to photoreceptor cell death (Paquet-Durand et al. 2009; Farber and Lolley 1974; Lolley et al. 1977), highlighting PKG as a target for the treatment of RD-type diseases.

In an effort to develop new drugs for the treatment of RD, a number of cGMP analogues designed to inhibit PKG have been synthetized (Butt, Eigenthaler, and Genieser 1994; Vighi, Trifunovic, et al. 2018). These compounds bear an Rp-configured phosphorothioate, which enables them to antagonize the activation of PKG by binding to the cGMP binding sites in the regulatory domain, without liberating the catalytic domain (Zhao et al. 1997). A previous study showed that the *in vivo* treatment with the cGMP analogue CN03, rescued photoreceptor viability and function in the genetically distinct *rd1*, *rd2*, and *rd10* animal models (Vighi, Trifunovic, et al. 2018). These results confirmed cGMP/PKG-signaling as a common target for the mutation-independent treatment of different RD-type diseases. Since then, several promising and potent 2^nd^ generation, novel cGMP analogues have been developed, but their biochemical properties and neuroprotective efficacy have not been examined thus far.

Here, we identified a novel cGMP analogue with strong photoreceptor-protective effects in retinal explants derived from *rd1* and *rd10* mice. We investigated the effects of this compound on retinal function in long-term organotypic retinal explant cultures derived from *rd10* and wild-type (WT) mice, using micro-electrode arrays (MEAs). The recordings revealed a stronger photoreceptor response and, surprisingly, also increased retinal ganglion cell (RGC) activity in the treated samples. Increased RGC viability in treated specimens was confirmed by histological analysis. Together, these results give new insights into the properties of cGMP analogues and other neuronal cell types.

## Materials and Methods

### Animals

C3H *Pde6b^rd1/rd1^* (*rd1*), congenic C3H wild-type (C3H), C57BL/6J wild-type (C57) and C57BL/6J *Pde6b^rd10/rd10^* (*rd10*) mice were housed under standard light conditions, had free access to food and water, and were used irrespective of gender. All procedures were performed in accordance with the ARVO declaration for the use of animals in ophthalmic and vision research and the law on animal protection issued by the German Federal Government (Tierschutzgesetz) and were approved by the institutional animal welfare office of the University of Tübingen.

### cGMP analogues synthesis

Synthesis of cyclic nucleotide analogues was performed by Biolog Life Science Institute GmbH & Co. KG according to previously described methods (Vighi, Trifunovic, et al. 2018) (https://patentscope.wipo.int/search/en/detail.jsf?docId=WO2018010965).

### *In vitro* PKG activation/inhibition assay

FLAG-Strep-Strep-tagged human PKG 1α (2–671) Wt, human PKG1β (4–686) Wt and human PKG2 (1-762) Wt were expressed in HEK293T cells. Cells were transfected at 80% confluency in whole medium employing the transfection reagent polyethyleneimine (Polysciences Europe GmbH, Germany). The cells were lysed using 50 mM Tris-HCl (pH 7.3), 150 mM NaCl, 0.5 mM TCEP, 0.4 % Tween, protease and phosphatase inhibitors (Roche, Germany). For purification we employed Strep-Tactin® Superflow® resin (IBA GmbH, Germany). We included an additional washing step with 366 mM Na2HPO4, 134 mM NaH_2_PO_4_ (pH 7.3) and 0.5 mM TCEP at room temperature to release any remaining nucleotides from the respective nucleotide binding pockets. Strep-tagged proteins were eluted with 200 mM Tris-HCl (pH 8), 300 mM NaCl, 2 mM EDTA and 5 mM desthiobiotin (IBA GmbH, Germany) and subsequently stored at 4°C in 50 mM Tris-HCl buffer (pH 7.3) containing 150 mM NaCl and 0.5 mM TCEP.

PKG kinase activity was assayed *in vitro* using a coupled spectrophotometric assay originally described by Cook et al. (Cook et al. 1982) in a clear 384 well PS-MICROPLATE (Greiner Bio-One, USA) in a CLARIOstar plate reader (BMG LABTECH, Germany). The final assay mixture contained 100 mM MOPS (pH 7.0), 10 mM MgCl2, 1 mM ATP, 1 mM phosphoenolpyruvate, 15.1 U/ml lactate dehydrogenase, 8.4 U/ml pyruvate kinase, 230 μM reduced nicotinamide adenine dinucleotide, 0.1 mg/ml BSA, mM β-mercaptoethanol and, as PKG substrate, 1 mM VASPtide (RRKVSKQE; GeneCust, Luxembourg). PKG Activation was determined with cGMP (Supplemental Figure S1) and the cGMP analogues CN003, CN226, and CN238 (all Biolog Life Science Institute GmbH & Co. KG, Germany) in dilution series ranging from 100 μM to 5.1 nM. The kinase reaction was started with 5 nM of the corresponding PKG isoform. Inhibition studies were performed by adding each PKG isoform supplemented with 2 μM cGMP to the assay mix and the respective cGMP analogue in dilutions ranging from 100 μM to 5.1 nM. One to three independent protein preparations were used for each assay, which in turn were performed in duplicates.

### Organotypic retinal explant cultures

Organotypic retinal cultures derived from C57, *rd10*, *rd1*, and C3H animals, were prepared as previously described (Belhadj et al. 2020; Caffé et al. 2001) under sterile conditions. Post-natal day (P)5 *rd1*, P9 or P12 *rd10* animals were sacrificed, the eyes rapidly enucleated and incubated in R16 retinal culture medium (07491252A; Gibco; Waltham, Massachusetts, USA) with 0.12% proteinase K (21935025; ICN Biomedicals Inc., Costa Mesa, California, USA) for 15 min at 37 °C. Proteinase K activity was blocked by the addition of 20% foetal bovine serum (FCS) (F7524, Sigma) followed by rinsing in R16 medium. Afterwards, the anterior segment, lens, vitreous, sclera, and choroids were removed, while the RPE remained attached to the retina. The explant was cut into a four-wedged shape resembling a clover leaf and transferred to a culture membrane insert (3412; Corning Life Sciences) with the RPE facing the membrane. The membrane inserts were placed into six-well culture plates and incubated with complete R16 medium with supplements and free of serum and antibiotics (Belhadj et al. 2020), in a humidified incubator (5% CO2) at 37 °C. For the first 48h the retinae were cultured with complete R16 medium without any treatment to allow adaptation to culture conditions. Afterwards, they were either exposed to different cGMP analogues (dissolved in water), each at [50 μM], or kept as untreated control. In both cases, medium was changed every second day with replacement of the full volume of the complete R16 medium, 1mL per dish, with fresh medium. The culturing paradigm was from P5 (explantation) to P11 (end of culture) for *rd1*. For the *rd10* model, two culturing paradigms were used: from P9 to either P17 or P19, and from P12 to P24. Culturing was stopped by 45 min fixation in 4% paraformaldehyde (PFA), cryoprotected with graded sucrose solutions containing 10, 20, and 30% sucrose and then embedded in Tissue-Tek O.C.T. compound (Sakura Finetek Europe, Alphena and enRijn, Netherlands). Tissue sections of 12 μm were prepared using Thermo Scientific NX50 microtome (Thermo Scientific, Waltham, MA) and thaw-mounted onto Superfrost Plus glass slides (R. Langenbrinck, Emmendingen, Germany).

### Histology

For retinal cross-sectioning preparation, the eyes were marked nasally, and cornea, iris, lens, and vitreous were carefully removed. The eyecups were fixed in 4% PFA, cryoprotected in sucrose, and sectioned as above.

### TUNEL assay

The various cGMP analogues were tested for their effect on photoreceptor cell death, using terminal deoxynucleotidyl transferase dUTP nick end labeling (TUNEL) assay (Loo 2011) (Sigma-Aldrich *in situ* Cell Death Detection Kit, 11684795910, red fluorescence). DAPI contained in the mounting medium (Vectashield antifade mounting medium with DAPI; Vector Laboratories, Burlingame, CA, USA) was used as nuclear counterstain.

### Immunofluorescence

Immunostaining with primary antibody against rabbit RBPMS (1:500; Abcam, Cambridge, UK) was performed on 12 μm thick retinal explants cryosections by incubating at 4 °C overnight. Alexa Fluor 488 antibody was used as secondary antibody. Sections were mounted with Vectashield medium containing 4’,6-diamidino-2-phenylindole (DAPI, Vector).

### Microscopy and image processing

Images were captured using 7 Z-stacks with maximum intensity projection (MIP) on a Zeiss Axio Imager Z1 ApoTome Microscope MRm digital camera (Zeiss, Oberkochen, Germany) with a 20x APOCHROMAT objective. For more details about the characteristics of the filter sets for the fluorophores used see Table 1. For the quantifications of positively labelled cells, pictures were captured on at least six different areas of the retinal explant for at least four different animals for each genotype. Adobe Photoshop (CS5Adobe Systems Incorporated, San Jose, CA) was used for image processing.

**Table 1:** Fluorophores and microscope filters. Excitation (exc.) and emission (em.) characteristics of the TMR red and AF488 and of the microscope filter sets used to visualize them.

| Fluorophore | exc. max. / em. max. (nm) | exc. filter / em. filter (nm) |
| --- | --- | --- |
| TMR red | 540/580 | 538-562/570-640 |
| AF488 | 495/519 | 450–490 /500-550 |

### *Ex-vivo* retinal function test

Prior to recording organotypic retinal cultures were kept dark for at least 12h, further manipulations were performed under dim red-light. The retinas were divided into two equal halves, one of which was placed immediately on the electrode field of the recording chamber and kept in the dark. The second retinal half was used for histological preparation and immunofluorescence (see above). Two recordings were obtained from locations within the central retinal half. Retinal function tests were performed in R16 medium, and the recording chamber temperature was set to 37°C. To record the light-evoked retinal responses, a micro-electrode array system (MEA; USB-MEA60-Up-BC-System-E, Multi Channel Systems; MCS; Reutlingen, Germany), equipped with HexaMEA 40/10iR-ITO-pr (60 electrodes = 59 recording and one reference electrode) was employed. The recordings were performed at 25.000 Hz sampling rate to collect unfiltered raw data. The trigger synchronized operation of the light stimulation (LEDD1B T-Cube, Thorlabs, Bergkirchen, Germany) and MEA-recording were controlled by a dedicated protocol implemented within the MC-Rack software (v 4.6.2, MCS) and the digital I/O – box (MCS). The light stimulation (white light LED, 2350 mW, MCWHD3, Thorlabs), guided by fiber-optic and optics, was applied from beneath the transparent glass MEA: five full field flashes of 500 ms duration with 20 s intervals. A spectrometer USB4000-UV-VIS-ES (Ocean Optics, Ostfildern, Germany) was employed to calibrate the intensity of the applied light stimulation (1,33E+14 photons/cm^2^/sec). For the analysis of the electrophysiology data, custom-developed scripts (MATLAB, The MathWorks, Natick, MA, USA) were used, if not indicated otherwise. MEA-recording files were filtered employing the Butterworth 2^nd^-order (MC-Rack, MC) to extract retinal ganglion cell spikes (high pass 200 Hz) and field potentials (bandpass 2 – 40 Hz). The field potentials recorded by the MEA system are referred as micro-electroretinogram (μERG), and largely correspond to the human electroretinogram (ERG) as described by (Stett et al. 2003). The filtered data were converted to *.hdf files by MC DataManager (v1.6.1.0). Further data processing was performed in MATLAB (spike and field potential detection) as previously described (Haq et al. 2018; Haq, Dietter, and Zrenner 2018).

### Statistics

#### 1) Analysis of PKG activation/inhibition assay

data were analyzed using GraphPad Prism 8.0.1 (GraphPad Software, Inc, La Jolla, CA, USA). Activation (K_act_) and inhibition (IC_50_) data are presented as mean ± standard deviation (SD); *n* = at least 3 except CN226 (n=2).

#### 2) Analysis of retinal cell death

The total number of TUNEL positive cells in the defined area of the outer nuclear layer (ONL) were estimated by dividing the ONL area by the average area occupied by a cell (*i.e*., cell size). The number of positively labelled cells in the ONL was counted manually on pictures captured on at least six different areas of the retinal explant for at least four different animals for each genotype. Only cells showing a strong staining of the photoreceptor nuclei were considered as positively labelled. Values obtained are given as fraction of total cell number in ONL (*i.e*., as percentage) and expressed as mean ± SD.

#### 3) Analysis of ganglion cell survival

An area of 1 mm^2^ was divided by the product of the length of counting and the section’s thickness (12 μm). The number of RBPMS positive cells was counted manually. For statistical analysis in both 1) and 2) a one-way ANOVA testing followed by the Dunnett’s multiple comparison test as implemented in Prism 8 for Windows (GraphPad Software) was conducted.

#### 4) Analysis of electrophysiology data

For the quantification of the PKG inhibitor effects on retinal light sensitivity, the photoreceptor (μERG) and ganglion cell (spikes) responses were considered: (1) Light responsiveness: This is represented by the percentage of light-dependent μERG detecting electrodes, to estimate the retinal light-sensitivity and to indirectly infer the density of functional photoreceptors in a given electrodes recording field. Note that a single MEA electrode captures the integrated signal of multiple photoreceptors within the recording field. The 59 MEA electrodes with 40 μm spacing together span an overall recording field of 340 × 280 μm, allowing to estimate the retinal light-sensitivity at 59 different positions. A μERG response upon light stimulation was counted as light-responsive if exceeding the respective threshold (response amplitude 1.75-fold ≥ calculated average of 500 ms control pre-stimulus baseline). (2) Deflection of the negative wave of the μERG: This measure reflects the strength of the light-evoked photoreceptor response – its hyperpolarization, equivalent to the a-wave in a conventional ERG –indicated by the initial negative deflection of the μERG (Figure 3 C, arrow). (3) Spike responses were accounted as light-stimulus correlated, if the post stimulus activity (average of 6 bin counts: 500 ms light duration and 100 ms post stimulation, 100 ms binning) exceeded the pre-stimulus activity (threshold: average of 5 bin counts pre-stimulus; 500 ms and 100 ms binning). Activity maps were generated to reflect the μERG recordings in their spatial context. Each pixel corresponds to a recording electrode of a MEA (center-center) and its surrounding recording area. The color encodes the negative deflection of the recorded μERG (−5 μV binning). For statistical analysis of the *rd10* data, one-way ANOVA followed by the Dunnett’s multiple comparison test was applied and for the WT dataset the Wilcoxon-Mann-Whitney test was utilized (MATLAB, The MathWorks).

## Results

### Novel PKG inhibitors protect *rd1* photoreceptors

Cyclic nucleotide (CN) analogues of cGMP (Figure 1) were previously generated in the context of the EU project DRUGSFORD (HEALTH-F2-2012-304963) and amongst others tested for their protective effects in primary rod-like cells. Here, we selected cGMP analogues that shared structural similarities with the retinoprotective compounds **CN003** and **CN004** (Vighi, Trifunovic, et al. 2018), which served as references. Structure wise all compounds feature a cGMPS backbone containing a sulfur-modified phosphate function with Rp-configuration, which confers PKG inhibitory properties (Zhao et al. 1997). Further modifications were introduced on the nucleobase moiety at positions 8 (R_1_) and 1, N^2^ (R_2_, R_3_). Therein the reference compounds contain a so-called PET-group (β- **p**henyl- 1, N^2^- **et**heno) at 1, N^2^ while this group is either lacking (**CN226**), substituted with an additional methyl-group (**CN238**), or replaced through the heteroaromatic furan ring (**CN007**). At position 8 the residue in **CN226** contains a phenyl ring as present in reference compound **CN004**, however deviating from **CN004** where a different, slightly larger linker has been introduced and said phenyl ring is unsubstituted. **CN007** and **CN238**, in turn, share the same bromide function as in the reference compound **CN003**. A compound concentration of 50 μM was used, based on previous *in vitro* results obtained with the reference compounds (Vighi, Trifunovic, et al. 2018).

**Figure 1:**
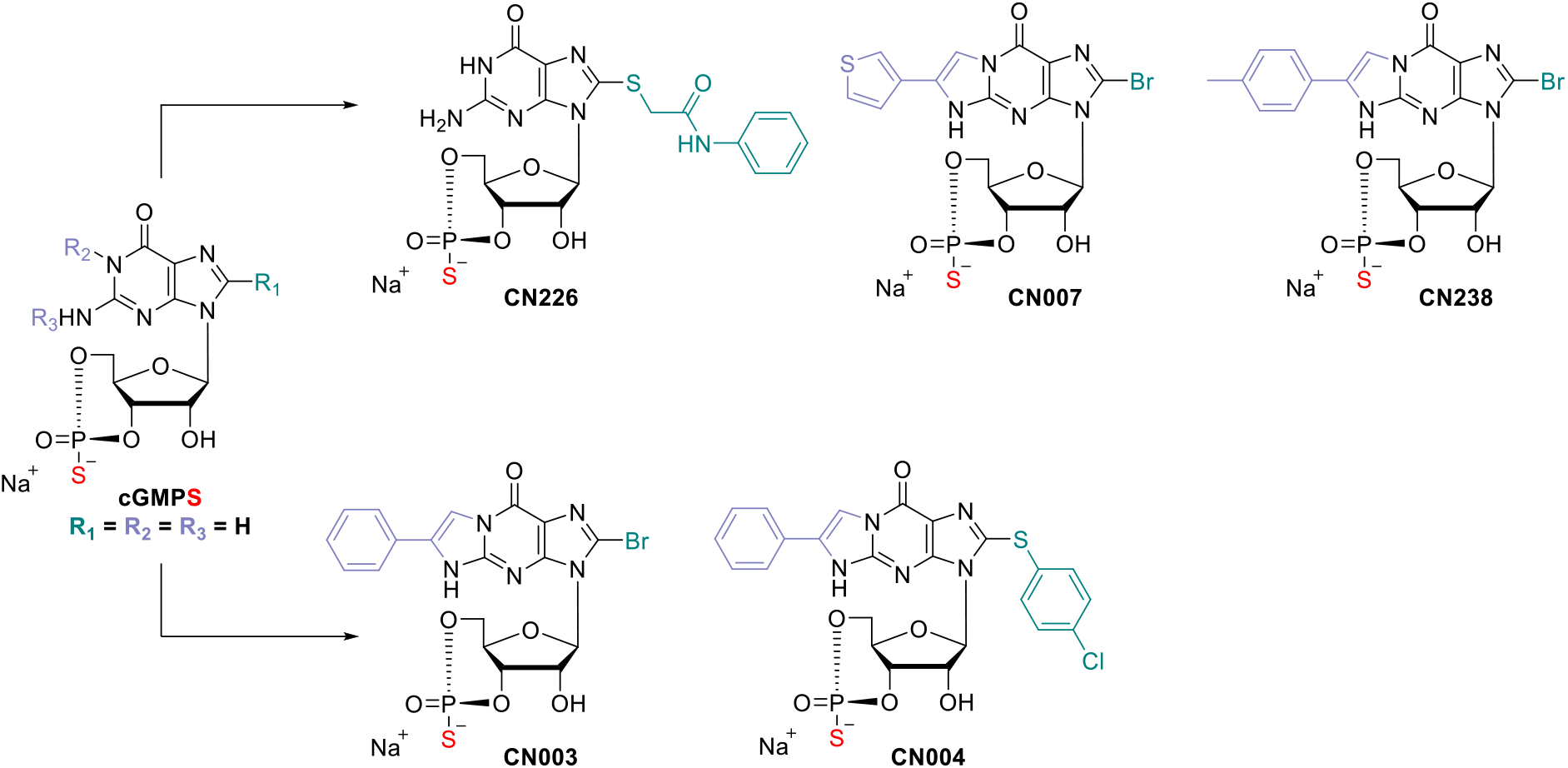
Structures of tested novel cyclic nucleotide analogues of cGMP and reference compounds. **CN003**: 8- Bromo- β- phenyl- 1, N^2^- ethenoguanosine- 3’, 5’- cyclic monophosphorothioate, Rp- isomer (Rp-8-Br-PET-cGMPS); **CN004**: 8- (4- Chlorophenylthio)- β- phenyl- 1, N^2^- ethenoguanosine- 3’, 5’- cyclic monophosphorothioate, Rp- isomer (Rp-8- pCPT-PET-cGMPS); **CN007**: 8- Bromo- (3- thiophen- yl- 1, N^2^- etheno)guanosine- 3’, 5’- cyclic monophosphorothioate, Rp- isomer (Rp-8-Br-(3-Tp)ET-cGMPS); **CN226**: 8-Phenylamidomethylthioguanosine- 3’, 5’- cyclic monophosphorothioate, Rp- isomer (Rp-8- PAmdMT-cGMPS); **CN238**: 8- Bromo- (4- methyl- β- phenyl- 1, N^2^- etheno) guanosine- 3’, 5’- cyclic monophosphorothioate, Rp- isomer (Rp-8-Br-pMe-PET-cGMPS).

We initially tested the cGMP analogues on organotypic retinal explant cultures derived from *rd1* mouse. The loss of function of PDE6 in *rd1* results in primary loss of rods already during development, with a peak around P13 and almost complete loss at P18 (Sahaboglu et al. 2013). Therefore, in *rd1* explants treatment with cGMP analogues started at P7 and ended at P11, *i.e*., at the onset of manifest retinal degeneration (Sancho-Pelluz et al. 2008), a time-point well suited for establishing possible protective effects.

As a readout of the effects of cGMP analogue treatments, we performed TUNEL assays on sections obtained from treated and non-treated (NT) specimens (Figure 2A). We considered any reduction in TUNEL positivity in the ONL indicative of a decrease in photoreceptor degeneration. When compared to *rd1* NT (100%), retinae treated with the compounds CN007, CN226, and CN238 (NT = 7.03 ±1.61; CN007 = 5.04 ± 1.26; CN226 = 6.18 ± 0.66; CN238 = 4.39 ± 0.50) showed a ≈32%, ≈17% and ≈40% reduction of TUNEL positive cells, respectively. The reference compounds CN003 and CN004 (CN003 = 4.79 ± 0.50; CN004 = 4.49 ± 0.45) confirmed their previously seen protective effects (Vighi, Trifunovic, et al. 2018) (Figure 2B).

**Figure 2:**
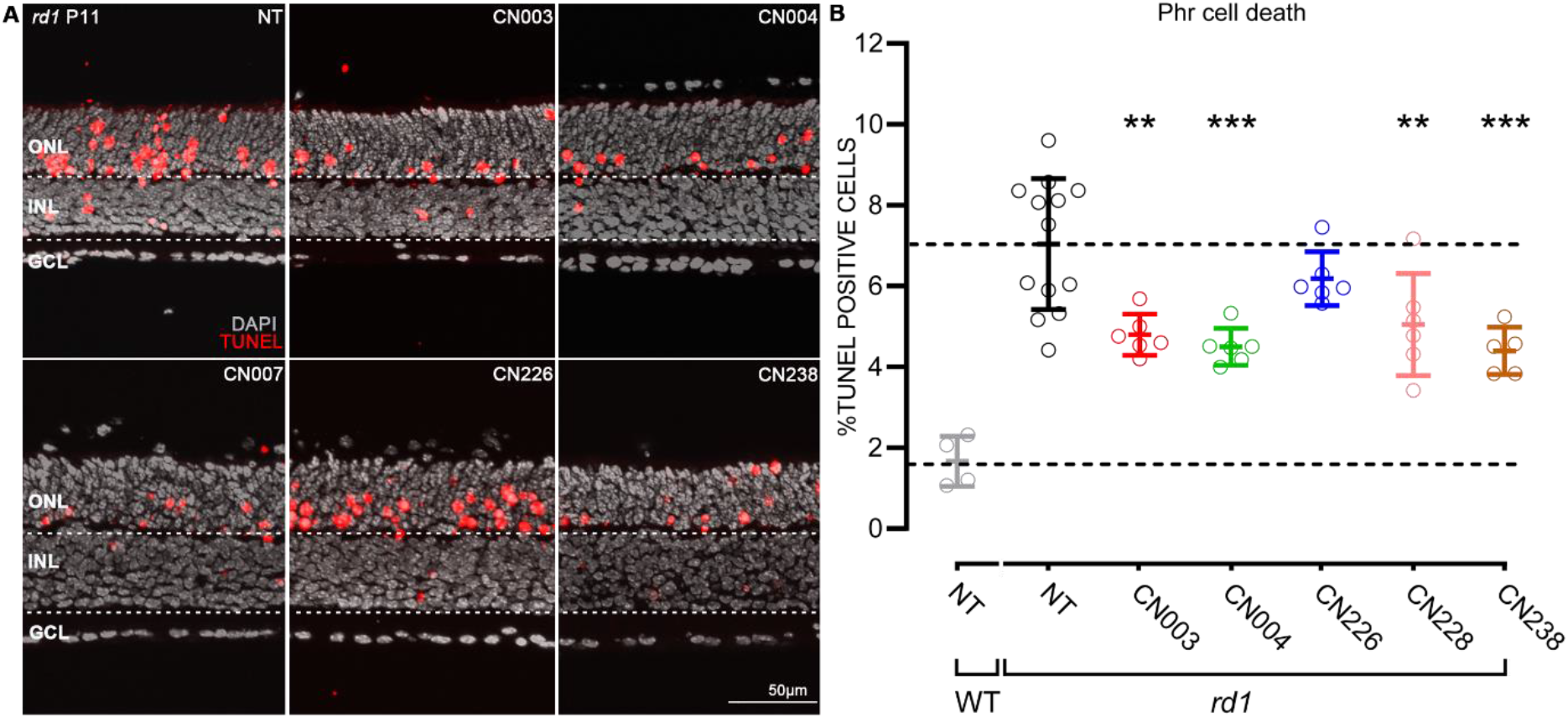
Retinoprotective effects of novel PKG inhibitors on *rd1* P11 organotypic retinal explant cultures. (**A**) Post-natal (P) day 11 *rd1* retinal explant cultures treated for four days with 50 μM of different cGMP analogues. TUNEL assay (red) indicated dying cells, DAPI (grey) was used as nuclear counterstain. (**B**) Quantification of TUNEL positive cells in the outer nuclear layer (ONL) of sections from A. Cell death rate in NT WT retina shown for comparison. A cell death rate lower than in the non-treated (NT) *rd1* retina was interpreted as evidence for photoreceptor protection. Compounds CN003 and CN004 had previously been established as photoreceptor protective (Vighi, Trifunovic, et al. 2018) and were used for reference. Testing was performed on n = 4 to 12 different retinae from different animals. Error bars: mean with SD. Statistical analysis was performed using one-way ANOVA followed by the Dunnett’s multiple comparison test; significance levels were: \*\**P* ≤ 0.01, \*\*\**P* ≤ 0.001. INL = inner nuclear layer, GCL = ganglion cell layer.

We then assessed the efficacy of the most promising compound CN238 to inhibit the PKG isoforms PKG1α, PKG1β, and PKG2. As references in this characterization, we included the previous lead compound CN003, as well as the compound CN226 as a “negative control” since it had not shown protection in *rd1* retinal explants (Supplemental Figure S2). This analysis indicated a slightly increased potency of CN238 towards PKG1 and PKG2 and furthermore revealed this compound to have partial agonistic effects on PKG1α at high concentrations. This suggests that CN238 is in fact modulating or dampening PKG1α activity rather than completely blocking it. On the other hand, analysis of the efficacy of CN226 showed that it was only weakly inhibiting PKG1β and PKG2, and that it did in fact activate PKG1α, in line with our results on *rd1* retinal explants (*cf*. Figure 1).

### PKG inhibition preserves viability and function of photoreceptors in *rd10* retinal explants

To assess the validity of these results across different animal models, we further tested the most promising compound CN238 and the reference compound CN003 on organotypic retinal explant cultures derived from *rd10* mice. As in *rd1* animals, the *rd10* mutation affects the gene encoding for the β-subunit of rod PDE6, however, in *rd10* photoreceptors the PDE6 enzyme retains some residual activity, delaying the onset of photoreceptor degeneration until P18 (Arango-Gonzalez et al. 2014; Power et al. 2020). Hence, the culturing and treatment paradigms were adjusted accordingly, and *rd10* retinae were cultured from either P9 to P17 or P19, or from P12 till P24. As in the *rd1* situation, and to allow the retinal explant to adapt to culture conditions, drug treatments were begun two days after explantation.

While the TUNEL assay gives a count of the number of dying cells at a given age, the number of mutant photoreceptors still surviving compared to the number of photoreceptors in an age-matched NT retina, gives the integral of cells preserved until a given time-point (Figure 3A). We therefore analyzed the number of photoreceptor rows to assess the long-term effects of the compounds. When CN003 and CN238 were tested on *rd10* retina in treatments reaching until P17 or P19 there was no statistically significant rescue, likely because of the comparatively late onset of *rd10* degeneration around P18. At P24, however, a clear and highly significant rescue of *rd10* photoreceptors was observed, with an increase in the photoreceptor row counts of ≈55% and ≈46% in samples treated with CN003 and CN238 (NT = 3.86 ± 1.01; CN003 = 6.01 ± 0.87; CN238 = 5.64 ± 0.32), respectively (Figure 3B).

**Figure 3:**
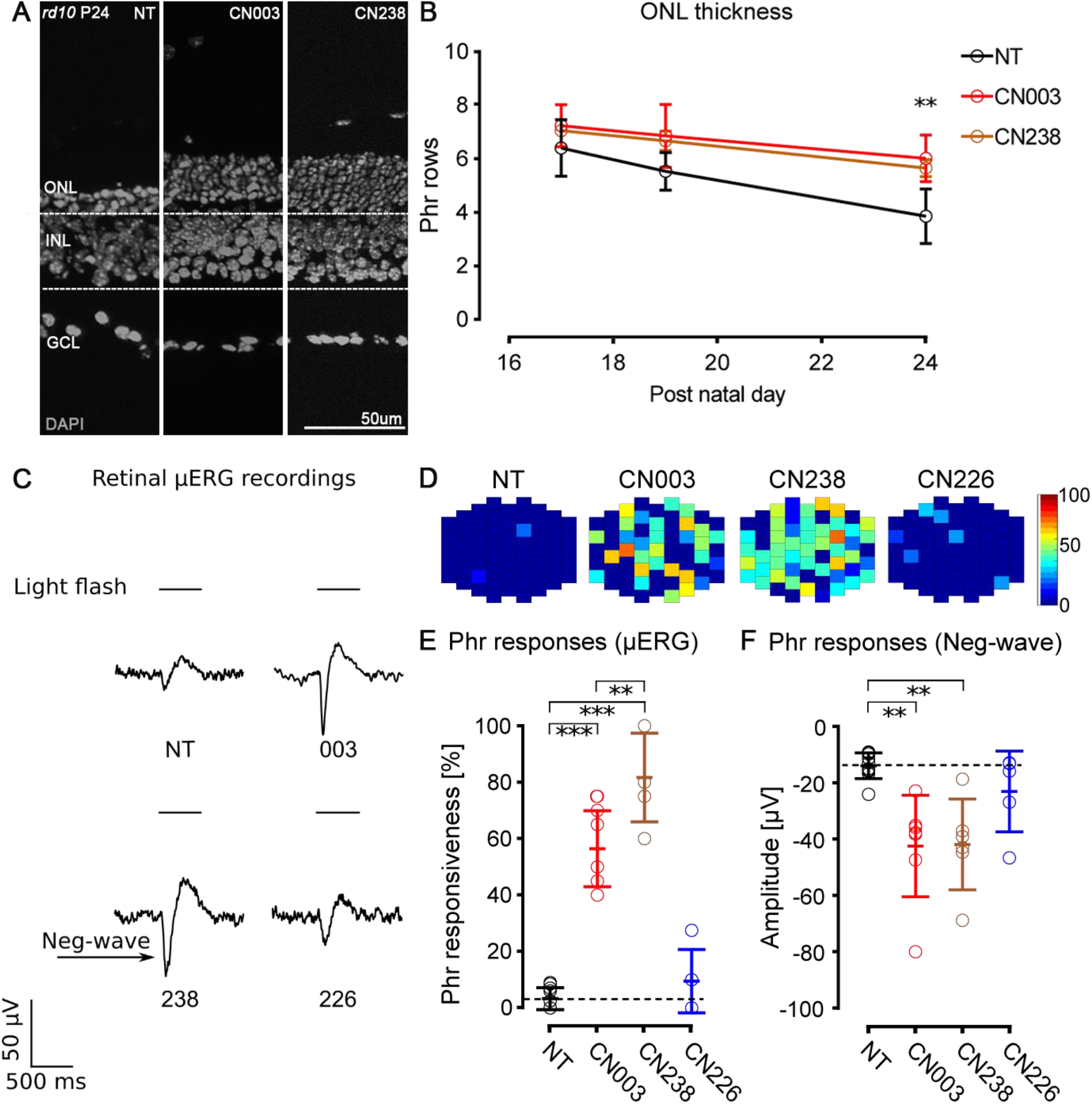
CN238 preserves photoreceptor viability and function in *rd10* retina: Organotypic retinal explant cultures derived from *rd10* mice were treated with cGMP analogues at 50 μM concentration. (**A**) Representative sections of post-natal (P) day 24 *rd10* retinal cultures treated or non-treated (NT) and stained with DAPI (grey). (**B**) *rd10* retinal explants were treated for varying times until either P17, P19, or P24, and photoreceptor rows were quantified. Row counts above NT were interpreted as evidence for photoreceptor protection. Testing was performed on n = 5 different retinae from different animals. (**C**) Representative μERG traces of NT and treated *rd10* retinal explants. The single-electrode data represents the integrated signal of multiple photoreceptors above a given electrodes recording field. The strength of the light-evoked photoreceptor hyperpolarization is indicated by the initial negative deflection (neg.-wave) of the μERG (arrow). (**D**) Activity map of representative μERG recordings across the 59 electrodes of the MEA, indicating the reactivity to light in spatial context. Each pixel corresponds to a recording electrode and the color from blue to red (*i.e*., 0 to 100) encodes the increasing intensity of the negative deflection of the recorded μERG. (**E**) Quantification of retinal light responsiveness as percentage of MEA electrodes displaying μERG negative deflections ≥ 1.75-fold average baseline. Testing was performed on n = 5 different retinae from different animals. (**F**) Average amplitudes of negative deflection in μERG recordings. Error bars: mean with SD. Statistical testing: one-way ANOVA with Dunnett’s multiple comparison test; significance level: \*\**P* ≤ 0.01. INL = inner nuclear layer, Phr = photoreceptor, μERG = micro-electroretinogram, GCL = ganglion cell layer, MEA = micro-electrode array.

To determine whether the increased photoreceptor survival seen with PKG inhibition also translated into improved retinal function, we performed electrophysiological recordings of *rd10* retinal explants employing a micro-electrode array (MEA) system and white light LED stimulator to apply 500 ms full-field light flashes. The method allowed for the selective assessment of photoreceptor functionality by recording light-elicited micro-electroretinograms (μERG). In this analysis, we included the compound that performed best in the retinal cell death assay, CN238 (*cf*. Figure 2), along with the previous lead compound CN003 and CN226, which here served as additional “negative” control.

As shown in representative μERG traces (Figure 3C), after 12 days of *in vitro* culture the PKG inhibitors CN003 and CN238 strongly increased the amplitudes of light-induced retinal responses, when compared to NT and CN226. In particular, the initial negative deflection of the μERG (Figure 3C, arrow), indicated a light-induced hyperpolarization of photoreceptors (*i.e*., the a-wave in a conventional ERG) and thus their functional preservation with drug treatment. For each retina, recordings were obtained from two different areas located dorsally and ventrally from the center and averaged (each recording area: 340 × 280 μm; 59 electrodes at 40 μm spacing). Since each of the MEA electrode captured the integrated signal of multiple photoreceptors within an electrode’s recording range, the total span of the MEA recording field allowed an estimation of the overall light-sensitivity of a given retinal explant. We considered a light-induced negative μERG deflection exceeding respective threshold (1.75-fold ≥ calculated average control baseline) to indicate light-responsiveness and drew activity maps for the MEA electrodes showing light responses (Figure 3D). We then expressed the number of electrodes showing such a response as percent of the total (Figure 3E). This analysis revealed that NT retina, and CN226 treated retina, displayed almost no response to light (NT = 3.3 ± 3.9 %; CN226 = 9.5 ± 11.2 %), while CN003 (56.4 ± 13.5 %), and even more so CN238 (81.7 ± 15.7 %), strongly and significantly increased light responsiveness of treated retinal explants (Figure 3E). In other words, treatment with CN003 or CN238 dramatically increased the retinal area responding to light by ≈ 17- or ≈ 25-times, respectively, when compared to NT. Compared to CN003, CN238 increased retinal responsiveness by yet another ≈ 45%, suggesting a further improvement of photoreceptor function and/or the density of the photoreceptors activating a given MEA electrode.

As an additional measure of photoreceptor functionality, we quantified the amplitudes of the initial negative μERG deflection, as a measure for the light-induced photoreceptor hyperpolarization. When compared to NT, retinal explants treated with CN003 or CN238 on average displayed a significantly stronger hyperpolarization response to light (NT: −13.6 ± 4.6 μV; CN003: −42.1 ± 18.0 μV; CN238: −41.5 ± 16.1 μV). In contrast, CN226 treatment had only a minor effect on response amplitudes (−22.7 ±14.3 μV) and/or the density of the photoreceptors activating a given MEA electrode (Figure 3F).

Taken together, we have established that the PKG inhibitors CN003 and CN238 preserved not only the viability of photoreceptor cells but also their functionality. In these comparisons the effects of the novel compound CN238 were at least equal, if not superior, to the previous lead compound CN003.

### CN238 prevents axotomy-induced degeneration of retinal ganglion cells (RGCs)

The MEA recordings of treated *rd10* retinal explants revealed another feature that was entirely unexpected: during the retinal explantation procedure the optic nerve is transected, and this axotomy leads to rapid degeneration and loss of most RGCs within 4-7 days of *in vitro* culture (Alarautalahti et al. 2019; Berkelaar et al. 1994; Osborne et al. 2016). Accordingly, in NT *rd10* retinal explants, cultured for 12 days *in vitro*, RGC spiking activity was virtually extinguished, and this was also true for CN226 treated explants. Yet, explants treated with CN003 or CN238 showed a very remarkable preservation of light-stimulus correlated RGC spiking activity (Figure 4A).

**Figure 4:**
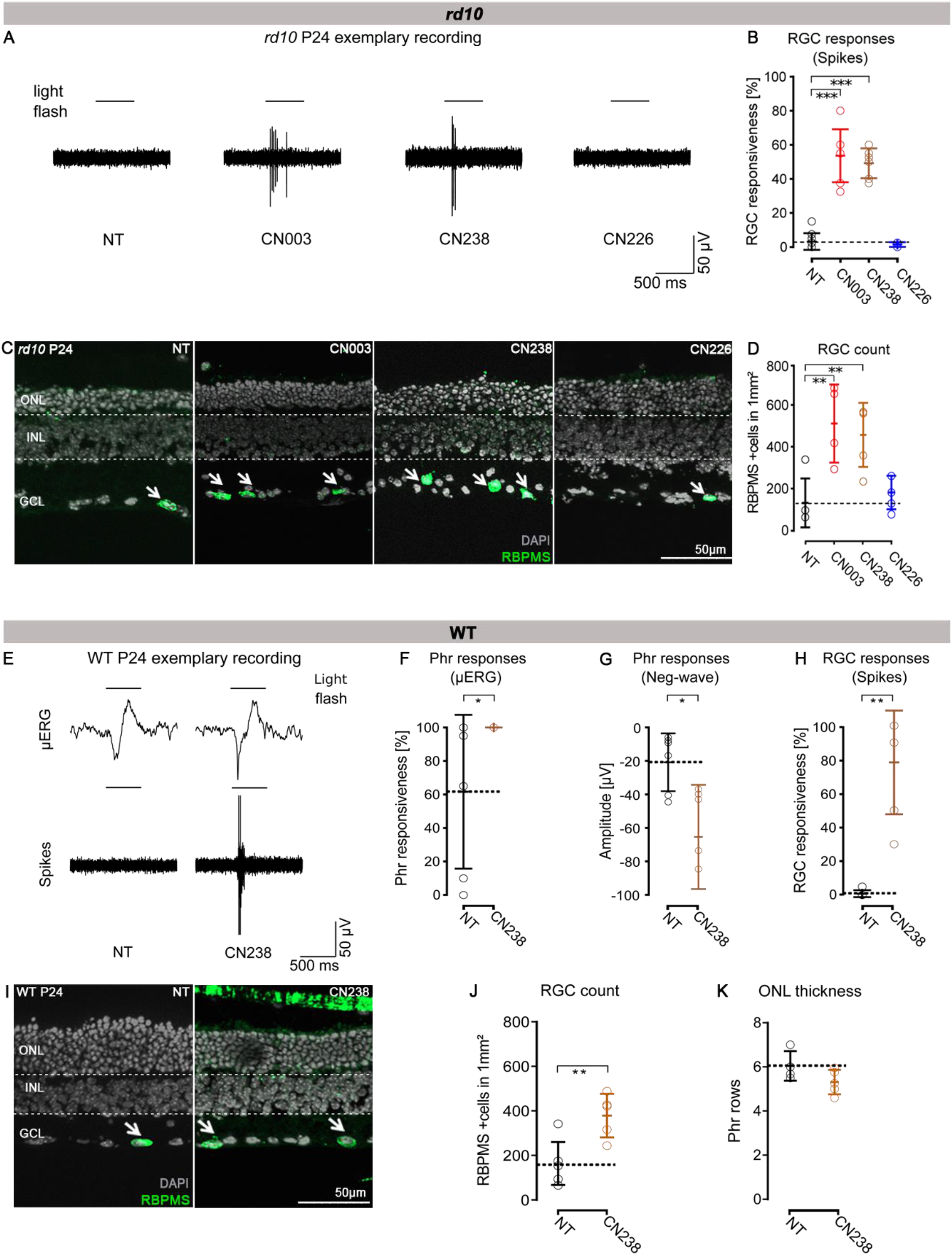
CN238 improves RGC viability and function in *rd10* and WT retinal explant cultures. (**A**) Representative light-correlated RGC spike MEA recordings performed on P24 *rd10* retinal explants. *rd10* retinas were treated from post-natal day (P) 14 to P24 with 50 μM of CN003, CN238, or CN226, and compared to non-treated (NT) retinal explants. (**B**) Quantification of light-evoked *rd10* RGC activity in NT and treated explants (percentage of MEA electrodes detecting light-correlated spike-activity). (**C**) Sections derived from recorded *rd10* retinal explant cultures were stained with DAPI (grey) and RBPMS (green). (**D**) Quantification of RBPMS positive cells in *rd10* P24 retinal explant sections. (**E**) Representative light-correlated RGC spike recordings of WT P24 retinal explants treated from P14 to P24 with 50 μM CN238 and compared to NT. (**F**-**H**) Quantification of light-stimulus evoked retinal activity in NT and treated WT explants: (**F**) light responsiveness (percentage of MEA electrodes detecting light-correlated μERG activity). (**G**) Quantification of negative μERG amplitudes. (**H**) RGC activity expressed as percentage of MEA electrodes detecting light-correlated spike-activity. (**I**) Sections derived from recorded WT P24 retinal explant cultures stained with DAPI (grey) and RBPMS (green). (**J**) Quantification of RBPMS positive cells in WT P24 retinal explants sections NT or treated with CN238. (**K**) Quantification of photoreceptor rows in NT and CN238 treated WT explants. Error bars indicate SD; statistical analysis in B, D: one-way ANOVA followed by Dunnett’s multiple comparison test; in F, G, H: unpaired Student’s *t*-test; levels of significance: \**P* ≤ 0.05, \*\**P* ≤ 0.01, \*\*\**P* ≤ 0.001. ONL = outer nuclear layer, INL = inner nuclear layer, Phr = photoreceptor, μERG = micro-electroretinogram, GCL = ganglion cell layer, MEA = micro-electrode array.

Similar to the analysis shown above for photoreceptor responses (μERG), we assessed the light responsiveness of RGCs across the entire surface of the MEA chip, from two different central recording areas. Here, light-induced RGC spike responses were considered to be stimulus correlated, if post stimulus activity (600 ms) exceeded the average of pre stimulus activity (500 ms, 100 ms bin, see methods for details). While RGC light responsiveness in NT and CN226 treated cultures was nearly absent (NT = 3.3 ± 4.9 %; CN226 = 1.5 ± 1.4 %), it was strongly and highly significantly increased in CN003 and CN238 treated retina (CN003 = 53.6 ± 15.5 %; CN238: 49.2 ± 8.8 %) (Figure 4B). Thus, in comparison to NT the area of the retina showing light responses at the RGC level was ≈ 16- or ≈ 15-times larger after CN003 or CN238 treatment, respectively.

We then used the very same retinal explants from which the MEA recordings were obtained for a histological workup, to assess the survival and physical presence of RGCs. To this end, we employed labelling for the RNA-binding-protein-with-multiple-splicing (RBPMS), a protein that is expressed in about 60% of RGCs (Rodriguez, de Sevilla Müller, and Brecha 2014) (Figure 4C). The quantification of RBPMS positive cells in the four experimental groups yielded very low RGC counts in NT and CN226 treated retina (NT = 134.1 ± 117.3 mm^2^; CN226 = 184.1 ± 81.06 mm^2^), while CN003 and CN238 treated explants displayed significantly larger numbers of RGCs (CN003 = 514.4 ± 187.7 mm^2^; CN238 = 460.3 ± 153.6 mm^2^). Compared to NT, retinal explants treated with CN003 and CN238 showed 3.8- and 3.4-fold higher numbers of RBPMS positive cells, respectively. In line with the previous experiments, the compound CN226 did not preserve the viability of RGCs (Figure 4D).

The magnitude of the rescue effect on axotomized RGCs raised the question whether this was a direct effect on RGCs or whether perhaps the preservation of *rd10* photoreceptors had indirectly enhanced the survival of *rd10* RGCs. To address this question, we extended our investigation to WT retinal explants, cultured for 12 days *in vitro*, from P12 to P24, treated or not with CN238.

The light-induced hyperpolarization response in the μERG obtained from P24 WT explants did not seem to differ between NT and CN238 treated retina, however, the RGC spiking activity was virtually absent in NT, but present after CN238 treatment (Figure 4E). The overall photoreceptor activity was obviously higher in WT than *rd10* explants, yet even in the WT situation CN238 treatment improved light responsiveness somewhat (NT = 61.7 ± 45.9 %; CN238 = 98.9 ± 2.6 %) (Figure F). In line with this, the light-response amplitude in terms of negative deflection of the μERG was also increased in CN238 treated explants compared to NT (NT = −26.2 ± 20.5 μV; CN238 = −79.0 ± 36.9 μV) (Figure G). The most striking effect of CN238 on WT retina was, however, observed at the level of RGC light-responsiveness: While NT retina showed nearly no light correlated activity, CN238 treatment largely preserved RGC light-induced spiking activity across the whole retinal explant (NT = 0.8 ± 2.0 %; CN238 = 78.3 ± 30.6 %) (Figure 4H). In relative terms CN238 had thus increased WT RGC functionality by a striking ≈ 95-times.

Further confirmation came from an investigation of WT RGC viability with RBPMS staining, which showed RGCs survival 2.3-fold higher in retinae treated with CN238 (NT = 163.9 ± 96.4 mm^2^; CN238 = 379.3 ± 97.7 mm^2^) (Figure 4J). In addition, we assessed photoreceptor survival in NT and CN238 treated samples, without detecting significant photoreceptor loss (Figure 4K).

These findings thus confirmed the formidable capacity of CN238 to preserve RGC activity despite the axotomy caused by the explantation procedure. This data also demonstrated that the RGC protection was independent of photoreceptor degeneration.

## Discussion

The excessive accumulation of cGMP in photoreceptors has long since been established as a trigger for the loss of photoreceptors in rare, RD-type diseases (Farber and Lolley 1974; Lolley et al. 1977). Here, using the novel inhibitory cGMP-analogue CN238, we validate PKG as the critical effector of cGMP-dependent cell death. Unexpectedly, the protective effect of PKG inhibition extended beyond photoreceptors to axotomized retinal ganglion cells, neurons whose degeneration is underlying common retinal diseases such as glaucoma and diabetic retinopathy. Importantly, a drug-mediated rescue of axotomized RGCs of the magnitude seen in this study has not been reported before. This emphasizes the general importance of PKG for neuronal cell death and highlights PKG inhibition as a new therapeutic approach for the treatment of neurodegenerative diseases in general.

### cGMP analogues as PKG inhibitors

Analogues of cGMP carrying an Rp-configurated phosphorothioate modification were first described as exceptionally potent and selective PKG inhibitors in the early 1990s (Butt, Eigenthaler, and Genieser 1994; Butt et al. 1995). While clinically used kinase inhibitors typically block the ATP-binding site present on all kinases (Atkinson et al. 2021; Johnson 2009), cGMP analogues target the cGMP-binding site present only on PKG, *i.e*., its physiological activation mechanism, thereby affording an extraordinary selectivity for PKG.

With CN238, we identified a cGMP analogue PKG inhibitor that preserved photoreceptor viability and function in *rd1* and *rd10* retina. This novel compound was found to have improved potency when compared to the reference compound CN003, a known PKG inhibitor with protective effects in *rd1*, *rd2*, and *rd10* mice *in vivo* (Vighi, Trifunovic, et al. 2018).

Both cGMP analogues are characterized by the ß-phenyl-1, N^2^-etheno (PET) group (Wei et al. 1996) and differ only by the additional methyl group in CN238. Interestingly, the compound CN007 which carried a similar N^2^-etheno modification was moderately photoreceptor protective, while CN226, which lacked such a R_2_-R_3_ modification (Figure 1), did not afford photoreceptor protection in *rd1* retina. This lack of efficacy was corroborated by the studies on *rd10* retina where CN226 showed significantly less preservation of photoreceptor function than CN003 or CN238.

The PET-group enhances the lipophilicity of Rp-cGMPS analogues, making it easier for the compounds to reach their target site inside the cell. In addition, the PET-group may bestow the ability to inhibit cyclic nucleotide gated ion (CNG) channels, albeit with an efficacy that is ≈ 2-3 log units lower than for PKG inhibition (Wei et al. 1996). Still, the protective effects of CN003 and CN238 could potentially stem from both PKG and CNG-channel inhibition. Indeed, CNG channel activity was for many years considered to be a driver of photoreceptor degeneration (Fox, Poblenz, and He 1999; Paquet-Durand et al. 2011). However, numerous studies in the last two decades explored the use of CNG-channel blockers, essentially without tangible results. Moreover, a recent study found that the selective block of CNG-channels with L-*cis*-diltiazem increased rather than prevented photoreceptor cell death (Das et al. 2020). Together, this makes it seem unlikely that the protective effects of CN003 / CN238 were due to CNG-channel inhibition. Nevertheless, a modulation of also CNG-channel activity by these compounds cannot be entirely excluded at this point.

### Effect of PKG inhibitors on photoreceptor function

The PKG inhibitors CN003 and CN238 robustly preserved the function of *rd10*-mutant photoreceptors as assessed via retinal recording of MEA μERG field-potentials. Each of the 59 MEA electrodes covered an area of approx. 80 μm^2^, meaning that the field potentials recorded likely originated from thousands of photoreceptors (Stett et al. 2003). Likewise, the amplitude of the initial negative deflection in the μERG represents the sum response of a large number of photoreceptors in a given recording field. Moreover, the recording of μERGs at different retinal locations allowed the creation of spatial activity maps for light responsiveness, both for treated and untreated tissues. The comparison of such maps of *rd10* retina revealed large differences between CN003/CN238 treated and untreated specimens, not only in terms of amplitudes of negative μERG deflections but also in terms of the areas of the retina showing responses to light flashes. This in turn demonstrates the magnitude of the photoreceptor protection over a large retinal area, an effect that was corroborated by the histological examination of the retina.

### PKG inhibition affords multilevel neuronal protection

The deleterious effects of high cGMP on photoreceptor viability were established already in the 1970s (Farber and Lolley 1974; Lolley et al. 1977). Yet, the role of PKG as a necessary and sufficient mediator of cGMP-dependent photoreceptor cell death was recognized only more recently (Paquet-Durand et al. 2009; Power et al. 2020). Accordingly, a systematic screening of PKG targeting cGMP analogues in various *in vitro* and *in vivo* models identified CN003 as a compound that afforded strong functional protection of *rd1*-, *rd2*-, and *rd10*-mutant photoreceptors (Vighi, Trifunović, et al. 2018).

The finding that PKG inhibition with either CN003 or CN238, in addition to photoreceptor protection, also prevented the demise of RGCs in both WT and *rd10* long-term retinal explant cultures was entirely unexpected. The transection of the optic nerve is a massive insult, known to cause rapid RGC function loss and degeneration (Berkelaar et al. 1994; Osborne et al. 2016). Instead, our MEA recordings indicated a striking preservation of RGC function, concomitant with a marked and significant increase in morphological RGC survival. A recent MEA study on retinal explant cultures found that in WT retina RGC activity gradually decreased to essentially zero within a culture period of 14 days. More importantly, RGC responses to light stimulation were no longer observed beyond 7 days of culture (Alarautalahti et al. 2019), a results that corresponds to our observations on both WT and *rd10* retina.

To investigate whether RGCs survival was somehow related to photoreceptor rescue, WT retinal explant cultures were treated with CN238 and compared with untreated specimens. Also, in treated WT retina the MEA recordings revealed strong RGC responses correlated to light stimuli. Immunohistochemical analysis, using the RGC marker RBPMS (Rodriguez, de Sevilla Müller, and Brecha 2014; Wang et al. 2016) and performed on the recorded retinal explants, demonstrated the strongly improved RGC survival after CN238 treatment. Remarkably, the number of RBPMS positive RGCs in both *rd10* and WT untreated retinal explants were approximately equal. Furthermore, in WT retina, the comparison of the number of photoreceptors in a row did not show any difference between the CN238-treated and untreated groups. These results suggest that in both the WT and the *rd10* situation, long-term retinal explant cultures display a loss of RGCs over time and that their rescue by CN238 is independent of photoreceptor survival. The failure of the cGMP analogue CN226 to preserve RGCs viability and function in *rd10* retinal explants indicates that RGC survival is connected to PKG inhibition.

The very marked RGC protection seen with CN003 and CN238 makes these compounds attractive for therapy development beyond photoreceptor diseases. Indeed, RGC degeneration is a hallmark of several retinal diseases with only limited treatment options to date. This includes glaucoma (Beykin et al. 2021), diabetic retinopathy (Lynch and Abràmoff 2017), exudative age-related macular degeneration (Medeiros and Curcio 2001), and non-exudative age-related macular degeneration (Yenice et al. 2015). How exactly PKG inhibition may afford RGC neuroprotection is not clear at present, yet numerous earlier studies have invoked detrimental effects of nitric oxide synthase and nitric oxide (NO) on RGCs (Mueller-Buehl et al. 2021; Neufeld, Sawada, and Becker 1999) also in optic nerve injury (Husain et al. 2014). Since, NO activates soluble guanylyl cyclase to produce cGMP and activate PKG (Bian and Murad 2014) increased NO production in injured RGCs will likely also cause overactivation of PKG, providing a rationale for the use of PKG inhibitors for RGC neuroprotection. The axotomy-induced degeneration of ganglion cells resembles a Wallerian-like retrograde degeneration (Howell et al. 2013; Vrabec and Levin 2007). The fact that PKG inhibition significantly reduces this type of degeneration thus indicates that PKG inhibitors may be applicable even more broadly in neurodegenerative conditions characterized by axonal damage, such as spinal cord injury or multiple sclerosis.

### Concluding remarks

While there has been tremendous progress in the development of new forms of therapy for RD, including gene, molecular and stem cell-based therapies (Hammond et al. 2021; Maguire et al. 2021), as well as retinal prostheses (Kitiratschky et al. 2015), there is still an important unmet medical need for more broadly applicable therapies that may benefit large groups of RD-patients. Our results support the idea that PKG/cGMP signaling is involved in photoreceptor degenerative processes (Power et al. 2020) and identify CN238 as a second-generation drug candidate with protective effects on photoreceptor survival and function in two mouse models for RD *in vitro*. However, further studies need to be conducted to assess whether the protective effects of CN238 can be extended to other models for RD characterized by abnormal cGMP signaling. In addition, the protective effect on RGCs in the explant culture system opens new perspectives for the use of PKG inhibitors for the treatment of common retinal diseases, including glaucoma.

## Conflict of interest declaration

A. R., F. S., and F.P.-D. have filed for three patents on the synthesis and use of cGMP analogues (PCTWO2016/146669A1, PCT/EP2017/066113, and PCT/EP2017/071859) and have obtained a European Medicine Agency orphan drug designation for the use of CN03 for the treatment of retinitis pigmentosa (EU/3/15/1462). F.P.-D. is shareholder of, or has other financial interest in, the company Mireca Medicines, which intends to forward clinical testing of cGMP analogues.

## Acknowledgements

We thank Norman Rieger for excellent technical assistance, Philipp Henning for his help in generating FSS-PKG constructs and Mathias Seeliger, Thomas Euler, Timm, Schubert, Per Ekström, and John Groten for helpful discussions. This research was funded by grants from the European Union (transMed; H2020-MSCA-765441), the Baden-Württemberg Foundation (BWST-WSF_006), the Charlotte and the Tistou Kerstan Foundation, and the German Ministry for Education and Research (BMBF; TargetRD, 16GW0267K, 16GW0269, 16GW0270).

## Supplementary information

**Figure S1:**
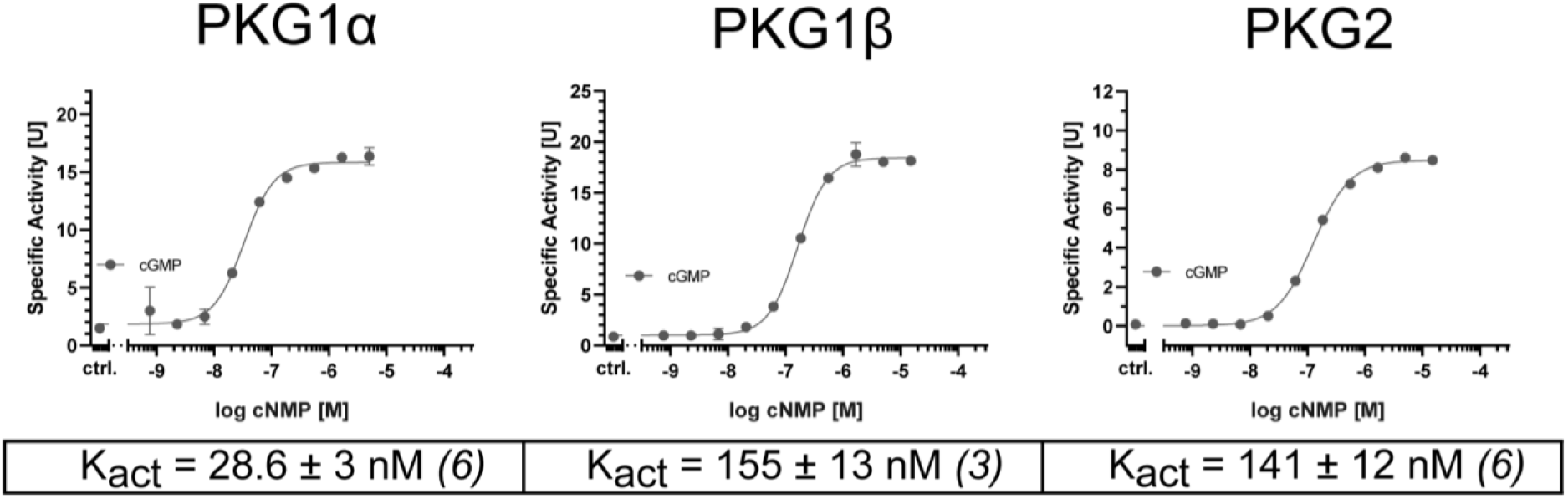
cGMP-dependent activation of the three distinct PKG isoforms. Activation curves of PKG1α, PKG1β, and PKG2 were obtained with cGMP dilution series ranging from 5 μM to 25.4 pm for PKG1α and from 15 μM to 762 pm for PKG1β and PKG2.

**Figure S2:**
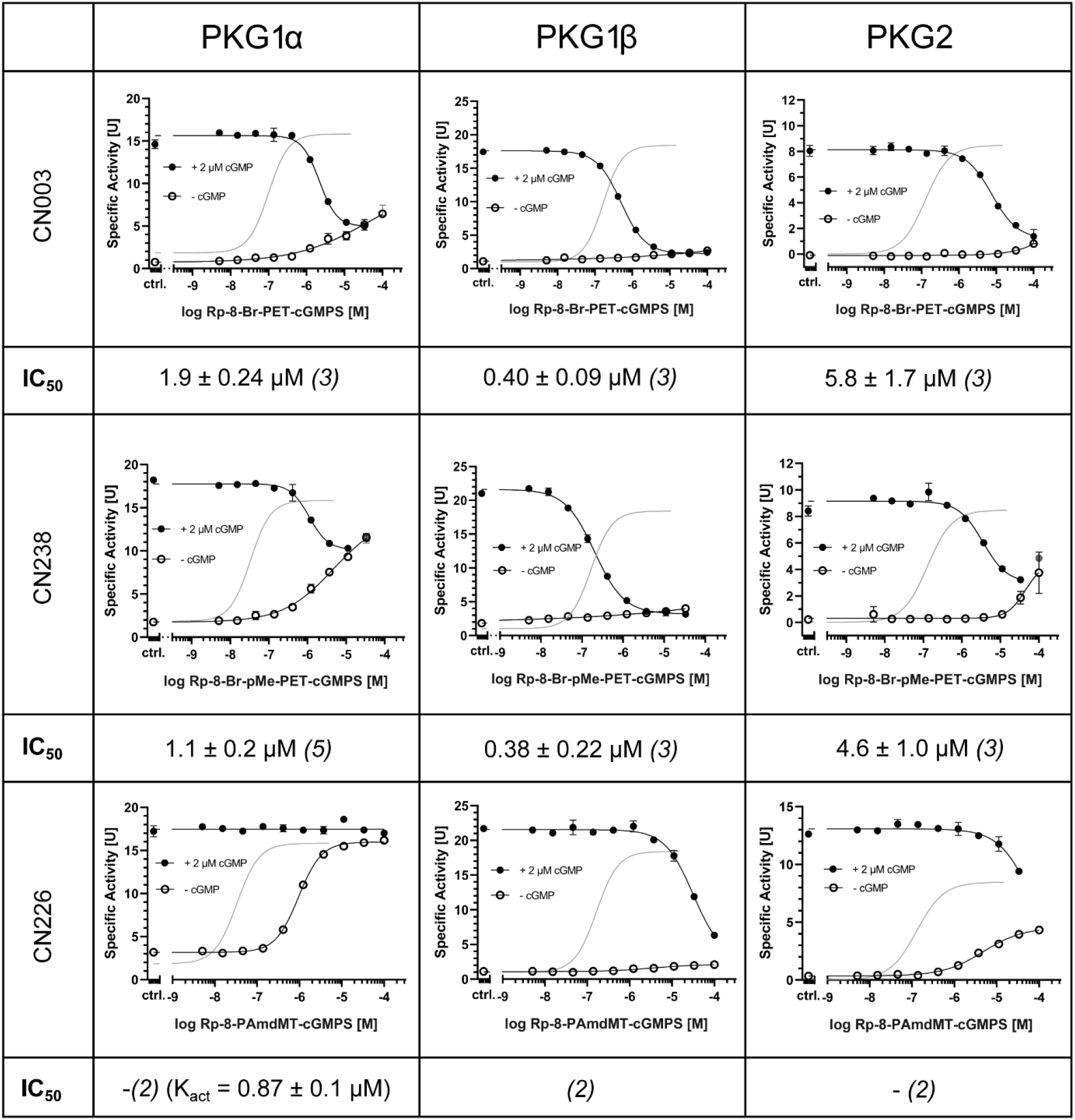
Agonistic and antagonistic properties of the cGMP analogues CN003, CN238, and CN226. The activation of all PKG isoforms (grey solid line) was determined with cGMP (*cf*. supplementary figure 1). Activation/Inhibition curves of 5 nM PKG1α, PKG1β, and PKG2 were measured with the cGMP analogues CN003, CN238, and CN226. For each analogue first the activation was determined (open circles) and in a second approach the inhibition in the presence of 2 μM cGMP was measured (solid circles). IC_50_ values were calculated from cGMP analogue dilution series ranging from 100 μM to 5.1 nM obtained in 2-5 independent measurements.

## Notes

### Summary of Updates

Acknowledgements and Authors conflict of interest update

